# Seg2Link: an efficient and versatile solution for semi-automatic cell segmentation in 3D image stacks

**DOI:** 10.1101/2022.10.10.511670

**Authors:** Chentao Wen, Mami Matsumoto, Masato Sawada, Kazunobu Sawamoto, Koutarou D Kimura

**Affiliations:** Graduate School of Science, Nagoya City University, Nagoya, Japan; RIKEN Center for Biosystems Dynamics Research, Kobe, Japan; Department of Developmental and Regenerative Neurobiology, Institute of Brain Science, Nagoya City University Graduate School of Medical Sciences, Nagoya, Japan; Division of Neural Development and Regeneration, National Institute for Physiological Sciences, Okazaki, Japan

## Abstract

Recent advances in microscopy techniques, especially in electron microscopy, are transforming biomedical studies by acquiring large quantities of high-precision 3D cell image stacks. However, to study cell morphology and connectivity in organs such as brains, scientists must first perform cell segmentation, which involves extracting individual cell regions of various shapes and sizes from a 3D image. This remains a great challenge because automatic cell segmentation can contain numerous errors, even with advanced deep learning methods. For biomedical research that requires cell segmentation in large 3D image stacks, an efficient semi-automated software solution is still needed. We created Seg2Link, which generates automatic segmentations based on deep learning predictions and allows users to quickly correct errors in the segmentation results. It can perform automatic instance segmentation of 2D cells in each slice, 3D cell linking across slices, and various manual corrections, in order to efficiently transform inaccurate deep learning predictions into accurate segmentation results. Seg2Link’s data structure and algorithms were also optimized to process 3D images with billions of voxels on a personal computer quickly. Thus, Seg2Link offers a simple and effective way for scientists to study cell morphology and connectivity in 3D image stacks.

## Introduction

In recent years, advances in microscopy techniques have allowed scientists to efficiently acquire large-scale/high-resolution 3D images with optical and electron microscopy (Hillman et al. 2019; Kornfeld et al. 2018; Xu et al. 2017). These large 3D images, particularly those obtained through electron microscopy, are useful for studying the structure and connectivity of cells in organs, such as brains (Parlakgül et al. 2022; Zheng et al. 2018; Wanner et al. 2016; Hildebrand et al. 2017; Lee et al. 2016). However, one of the necessary procedures, segmenting cells into individual ones, especially in 3D images, is still a challenging and time-consuming task and thus becomes a bottleneck (Lichtman et al. 2014; Kornfeld et al. 2020).

Many factors contribute to the difficulty of cell segmentation in 3D images, and three of the most important ones are as follows: 1) Cell boundaries can be difficult to identify in many biomedical images, so even a human annotator must infer them based on context and experience. Given that automatic segmentation techniques such as deep learning are still inferior to human experts, automatic segmentation results often contain a large number of incorrect cell boundary predictions, resulting in incorrect segmentation and requires manual corrections. 2) Slice displacement along the z-axis, particularly non-rigid displacement, is common in 3D cell images, again resulting in incorrect segmentation and requires manual corrections. 3) Recent 3D biomedical images, particularly those obtained by electron microscopy, typically have a large image size and a large number of cells, which substantially increases the processing time for automatic segmentation and manual corrections, as well as the requirement for large memory and disk space.

Traditionally, segmenting large 3D cell images requires time-consuming manual annotations (Cardona et al. 2010; Helmstaedter et al. 2011). Recent advances in machine learning, especially deep learning methods (Berning et al. 2015; Falk et al. 2019; Stringer et al. 2021; Januszewski et al. 2018), have made automatic cell detection and segmentation practical, yet existing automatic methods rarely achieve the accuracy of human experts and still requires intensive efforts to correct the errors manually. For example, currently one of the most advanced methods is called flood-filling networks (FFNs) (Januszewski et al. 2018), which requires users to prepare a large amount of manually annotated training data and is computationally expensive. Even with this method, automatic segmentation of real 3D brain images has a large number of errors that take a long time to correct (Scheffer et al. 2020). In addition, due to the complexity of 3D segmentation, even a local error in a specific slice in deep learning predictions can lead to a completely incorrect segmentation in a large 3D space, making handling the errors in automatic segmentation results a difficult task (Supplementary Fig. 1). Furthermore, developing a high-precision automatic segmentation method that is suitable for all conditions is difficult due to the complexity and variability of cell morphology and imaging conditions used by different research groups.

For the reasons stated above, we consider that automated segmentation methods will continue to require manual corrections to produce accurate results for the foreseeable future. Therefore, if we can develop a semi-automatic software program that combines automatic segmentation and manual correction while maximizing operational efficiency, it will substantially help scientists to quickly and accurately segment 3D images. The following features would be ideal in such a software program: 1) Be able to generate automatic segmentation using the imperfect cellular/non-cellular predictions from deep learning techniques. 2) Allows the user to correct errors in automatic segmentation on a slice-by-slice basis, and can utilize the corrected results to more accurately segment subsequent slices. 3) Possesses optimized procedures of manual corrections as well as optimized computational and storage efficiency, in order to reduce user operation time and hardware requirements. Previous software programs for assisting 3D cell segmentation could not utilize the cellular/non-cellular predictions as input and/or lack some important functions for improving efficiency (Helmstaedter et al. 2011; Cardona et al. 2012; Haehn et al. 2014; Berger et al. 2018; Zhao et al. 2018; Urakubo et al. 2019).

Here we developed Seg2Link, a semi-automatic segmentation software that takes imprecise cell boundary predictions based on deep learning as inputs, automatically performs 2D segmentation and cross-slice linking, and allows users to correct the segmentation results quickly. Seg2Link can segment entire images as well as user-defined regions. Seg2Link offers a number of functions, including cell merging, deletion, division, division + relink, cell sorting by size, fast cell localization, multi-step undo/redo, fast saving and reloading of intermediate segmentation results, and more, allowing users to easily check and quickly correct the segmentation results. The comparisons of functions in Seg2Link with those in other software programs are summarized in Table 1, and described in detail below. By optimizing the underlying data structures and algorithms, our software reduces computation time and increases storage efficiency, allowing users to segment 3D cell images with billions of voxels on an ordinary desktop or laptop computer. We believe Seg2Link successfully integrates many of the key functions required by semi-automatic 3D cell segmentation and can help scientists in analyzing the cell morphology and connectivity in organs such as brains more efficiently.

**Table 1.** Comparison of 3D segmentation software features. ✓: has the feature. △: partially has the feature. - : does not have the feature.

|  | Automatic segmentation |  |  | Manual Correction / Inspection |  |  |  |  |  |  |  | Multiple platform | Open source |
| --- | --- | --- | --- | --- | --- | --- | --- | --- | --- | --- | --- | --- | --- |
|  | Predict cell/noncell regions | Segment cell regions into individual cells | Mask | Merge | Delete | Division (2D) | Relink after Division | Division (3D) | Undo/ Redo | Sort by sizes / Remove tiny | Locate cells |  |  |
| Ilastik (Berg et al. 2019) | ✓ | ✓<br>watershed 3D | - | - | - | - | - | - | - | - | - | ✓ | ✓ |
| TrackMate (Ershov et al. 2021) | - | △<br>2D SEG (imported from other software) + Link | - | - | - | - | - | - | - | - | - | ✓ | ✓ |
| DoJo (Haehn et al. 2014) | - | - | - | ✓ | ✓ | ✓ | - | - | ✓ | - | - | △<br>not in Windows | ✓ |
| UNI-EM (Urakubo et al. 2019) | ✓ | ✓<br>watershed 3D | - | ✓ | ✓ | ✓ | - | - | ✓ | - | - | △<br>not in macOS | ✓ |
| NeuTu (Zhao et al. 2018) | - | - | - | ✓ | ✓ | - | - | ✓<br>requires adding seeds | △<br>cannot undo after confirming a merge | - | ✓ | △<br>not in Windows | ✓ |
| VAST Lite (Berger et al. 2018) | - | - | - | ✓ | ✓ | △<br>but only with straight line | - | - | - | - | ✓ | - | - |
| Seg2Link | - | ✓<br>watershed 2D + Link | ✓ | ✓ | ✓ | ✓ | ✓ | ✓<br>requires annotating boundary | ✓<br>multiple steps | ✓ | ✓ | ✓ | ✓ |

## Results

### Overview

Seg2Link is made up of two modules (Fig. 1A). The first module, Seg2D+Link, takes cellular/non-cellular predictions on each pixel generated by deep learning (or other techniques) as input and automatically segments the cell regions into individual cells in each slice, and then links the cells with previous slices along the *z*-axis (Fig. 1A, left and middle panels). Users can manually correct the segmentation results immediately after the automatic segmentation in each slice, so that to help the software improve the segmentation quality in subsequent slices. After all the slices have been segmented and corrected, the 3D segmentation result can be exported and imported into the second module, 3D Correction, which allows the user to comprehensively check and correct any remaining errors in each 3D-segmented cell (Fig. 1A, right panel). The final corrected 3D segmentation results can be exported as image sequence files for further analyses in other software.

**Figure 1.**
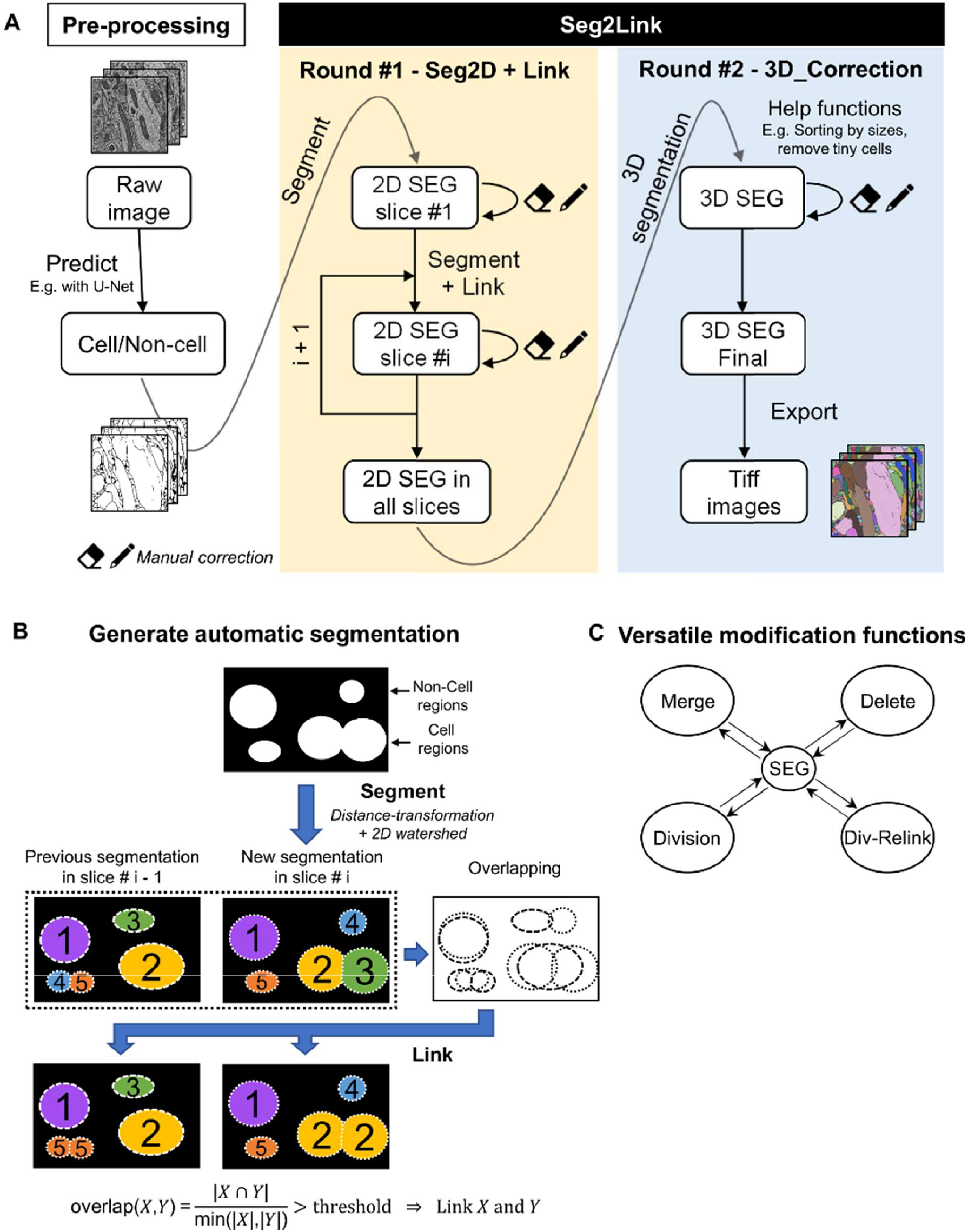
Seg2Link workflow and core functions. (A) Seg2Link workflow. (B) The methods for automatic segmentation. (C) The manual modification functions. SEG is an abbreviation for segmentation.

We created a set of graphical user interfaces (GUI) using the napari library (napari contributors, 2019) to allow users to specify images and parameters, and perform semi-automatic segmentation. Users can use the tool in napari to check and edit the labels in the results and can use our pre-defined hotkeys and GUI buttons to perform automatic segmentation and manual corrections (see next section).

### Semi-automatic segmentation

This software’s core module, Seg2D+Link, allows users to perform rapid slice-by-slice automatic segmentation and manual corrections. Users can use other programs to generate deep learning predictions (We provided one. See Methods), which can then be exported as TIFF images and used as inputs to the Seg2D+Link module. In the automatic segmentation part, the program applies a 2D watershed (Beucher and Meyer, 1993) to the cellular/non-cellular predictions by a pre-trained deep neural network and generates segmented cell regions in each slice (Fig. 1B). From the second slice, our program automatically links the segmented cells with cells in previous slices along the z axis using the overlap linking method (Fig. 1B). The software allows users to freely correct the errors in the automatic segmentation results using various commands (Fig. 1C). Users can also apply multi-step undo and redo functions to user operations to quickly go back to the previous states in case a misjudgment occurs, which is not a built-in function in the napari platform and is not supported by some of the 3D segmentation software. Our software also automatically saves the segmentation results in every slice to the hard disk, allowing the user to resume the previous results later. We also accelerated the modification and the caching/saving processes by designing a custom data structure (Supplementary Fig. 2). During our testing, the automatic segmentation + manual correction takes ~3 minutes per slice on the first 10 slices of the demo dataset with ~700 cells per slice, using a laptop computer (See Methods). Specifically, manually correcting the first slice (without automatic linking) took 8 mins, while manually correcting each subsequent slice (with automatic linking) from #2 took only ~2 mins, indicating a substantial increase in efficiency by performing automatic linking across slices.

The main window of the Seg2D+Link module displays the current segmentation results, as well as raw and prediction images in different layers (Fig. 2A). The automatic segmentation and the manual correction functions in Seg2D+Link were carefully designed so that users can easily perform the segmentation. First, 2D watershed + link typically produces good segmentation results with few mistakes. Although the segmentation in the first slice frequently contains a few over-segmented areas, they are easily corrected with the merge command described in Fig. 1C, and the manually performed merge will guide the program to automatically merge the over-segmented areas in the next slice using the overlap linking algorithm, which greatly improves the segmentation quality of the following slices (Supplementary Fig. 3). As a result, users typically only need to make few manual corrections in the slice #2 and afterwards (Fig. 2B). Additionally, merge/delete operations performed on any subsequent slice are also automatically applied to previous slices to improve operational efficiency. Secondly, Seg2Link’s correction function requires very few manual operations to perform. For example, relinking cells after division requires only editing the cell boundary in slice # i and then pressing the hotkey R to finish the division and relinking with the previous slice # i-1 (Fig. 2C). Merging or deleting cells only requires a mouse click and pressing the hotkey A to select each cell and add it to the list, followed by a press of M or D to complete the merging or deleting (Fig. 2C). Furthermore, when users are only interested in a specific subregion of a 3D image, Seg2Link allows them to specify the subregion with a mask image and segment the specified region selectively (Fig. 2D).

**Figure 2.**
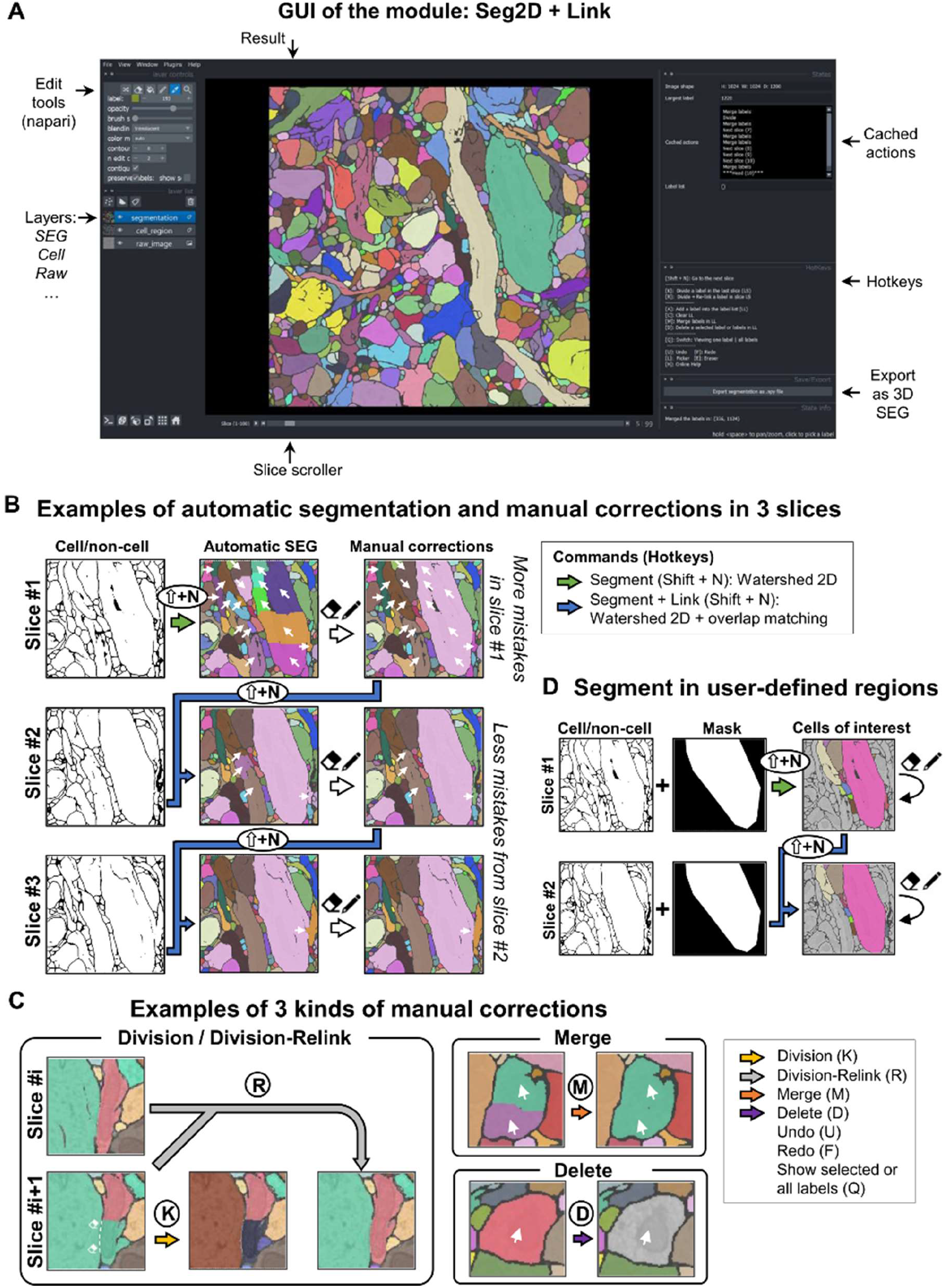
Functions of the Seg2D+Link module. (A) The GUI of the main window. (B) The process of semi-automatic segmentation in slices #1, #2, and #3. Different colors indicate different cells. The arrows on the images indicate the regions with errors in the automatic segmentation and the results after manual corrections. (C) The manual correction functions. Arrows on the images indicate the regions to be corrected. (D) The masking function for segmentation in user-defined regions.

### Comprehensive inspection and correction

The second module, 3D correction, allows the user to perform a comprehensive checking and correction of the 3D segmentation results made in the Seg2D+Link module (Fig. 3A). Since the majority of the errors have already been corrected in the Seg2D+Link module, we designed two functions to make it easier for users to inspect the segmentation results in the entire 3D image. First, large cells are potentially more important, but inspecting all large cells in a 3D image with thousands of cells is difficult. Our software allows users to sort 3D cells by volume size and remove cells that are smaller than a user-defined threshold (Fig. 3B, left), so that users can inspect the large cells sequentially by ID. Second, because a single cell typically occupies tens of slices, a small portion of all slices, visually searching for a cell with a specific ID among thousands of slices is time-consuming and boring. Our software allows users to quickly locate a specific cell and jump to the middle slice of it (Fig. 3B, right). With these two functions, users can quickly find and inspect large cells.

**Figure 3.**
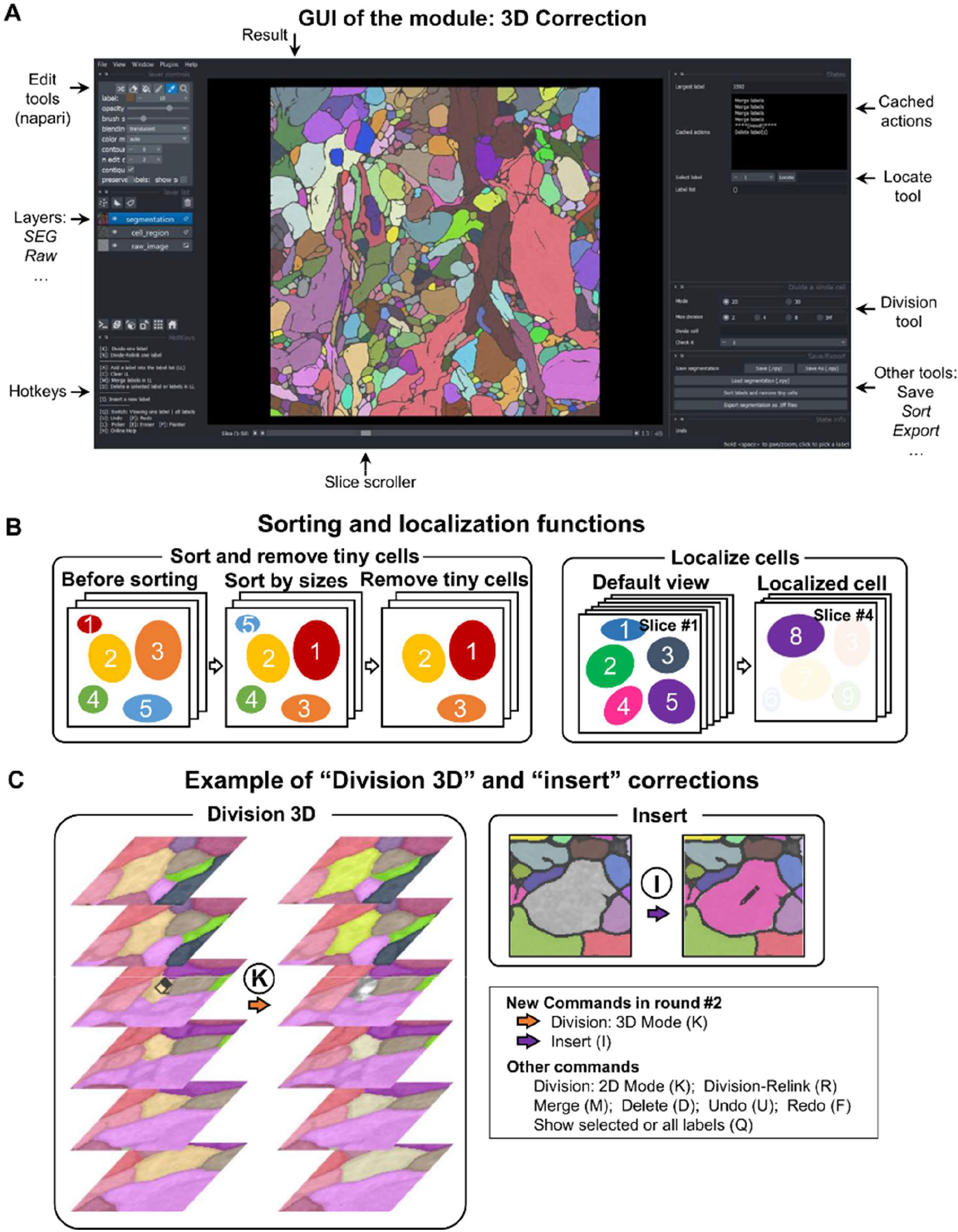
Functions of the 3D correction module. (A) The GUI of the main window. (B) The cell sorting, cell removal, and cell localization functions. (C) The manual correction functions. The eraser and pen on the images indicate the cleared cell region and the newly inserted cell region, respectively.

Aside from the two inspection functions mentioned above, we also added two manual correction functions that are needed for the 3D correction module but not appropriate to perform in the Seg2D+Link module. First, instead of dividing a cell in each 2D slice, users can divide it in 3D space, which is useful for separating two cells that are incorrectly linked along the z-axis (Fig. 3C, left). Second, users can insert new cells that have not been detected by deep learning predictions (Fig. 3C, right).

In addition, we improved the computational efficiency of cell localization, which is critical because all implementations of manual corrections require the cell to be localized first. Since the time required for localization grows with the 3D image size, this can become a serious problem. We speed up this function by storing each cell’s bounding box (bbox) information in a cache (Supplementary Fig. 4), so that the software can search for a cell in a much smaller subregion, and the time required for localization is primarily determined by cell size rather than the image size. During our tests on a larger dataset than the demo dataset (see Methods), we randomly selected 1/50 of the 52, 237 segmented cells and found that localizing a cell without using bbox takes 2.87 secs on average (5th and 95th percentile: [2.84, 2.92]), whereas localizing a cell using bbox takes 0.028 secs on average (5th and 95th percentile: [0.00, 0.09]), indicating a 103-times acceleration.

The default workflow for Seg2Link is to use Seg2D+Link as the first round to generate the segmentation and correct the majority of the errors, followed by 3D Correction as the second round to correct the remaining errors (Fig. 1A). Aside from that, Seg2Link also allows users to import 3D segmentation results obtained with 3D watershed from other software and then correct the errors with the 3D correction module. The second solution may appear to be more efficient because users can avoid the first round of manual correction. However, at least in large datasets, it is not the best choice for a variety of reasons. First, 3D Correction requires much more memory/storage and time than Seg2D+Link to cache/save intermediate results (Supplementary Fig. 5), because the software must store the entire 3D array rather than the 2D array + a small list as in the Seg2D+Link module, making it less efficient for correcting a large number of errors in the automatic segmentation results. Second, our results on the demo dataset show that 3D watershed can produce more boundary errors than 2D watershed + overlap linking (Fig. 4), likely because cell boundaries in each x-y slice are inferred with 3D watershed using boundaries in neighboring x-y slices along the z-axis. Correcting such incorrect boundaries requires users to manually paint many pixels, increasing the time cost. Finally, the 3D watershed is computationally expensive, requiring 0.037 GB RAM per slice in the demo dataset when tested with the MorphoLibJ plugin (Legland et al. 2016), whereas the Seg2D+Link and 3D correction modules require only 0.7 GB and 2.7 GB, respectively, for processing the entire demo dataset with 1,200 slices. For these reasons, we recommend users follow the default workflow (i.e., Seg2D+Link).

**Figure 4.**
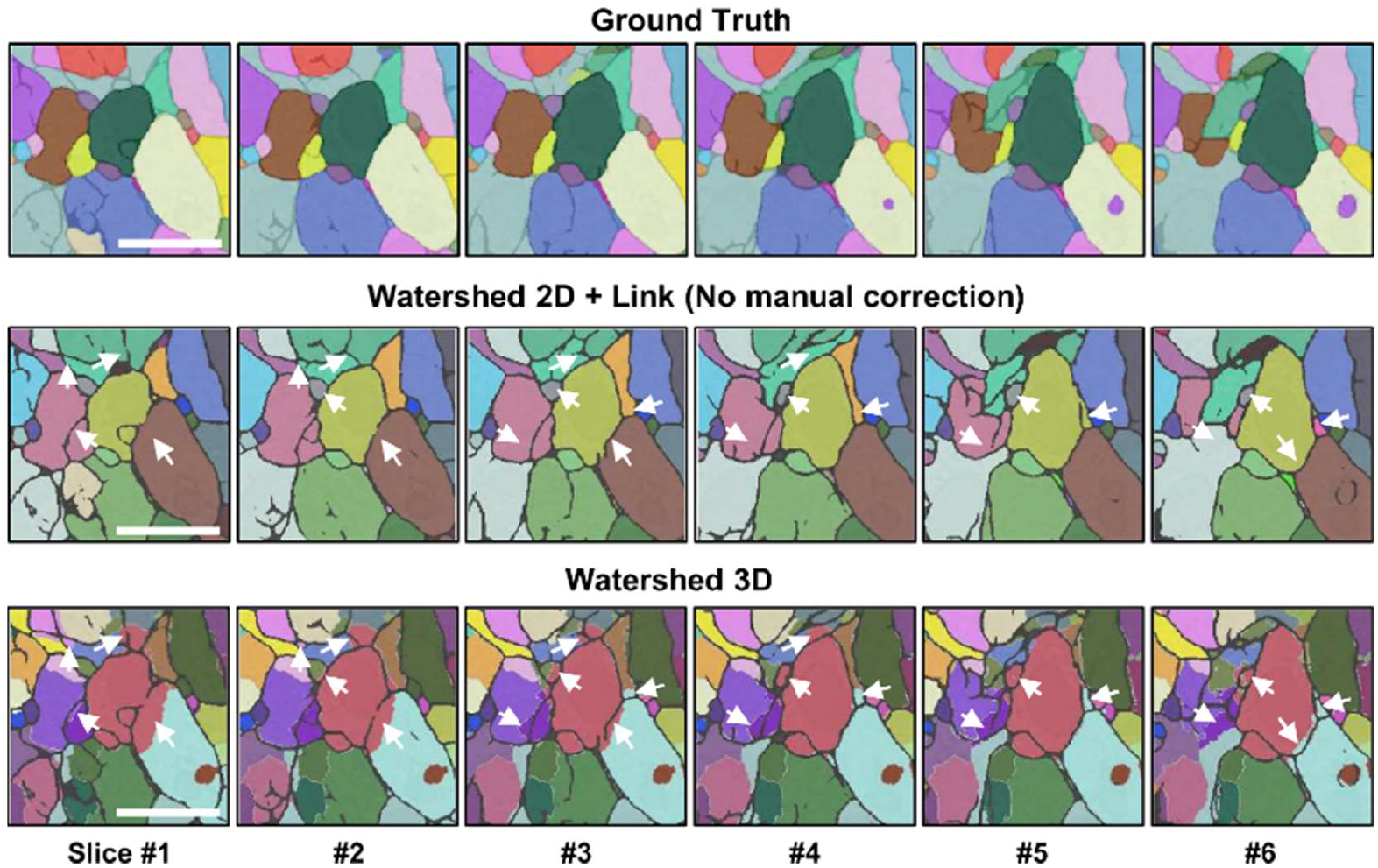
The segmentation results with two different approaches. (Top) The ground truth indicating the correct segmentation results. (Middle) Segment cells with 2D watershed + overlap linking using Seg2D+Link, without performing manual correction. (Bottom) Segment cells with 3D watershed using the “Distance Transform Watershed 3D” function in the plugin MorphoLibJ in ImageJ (Legland et al. 2016). Scale bars: 1 μm. The arrows indicate the same regions with correct (watershed 2D + link and ground truth) and incorrect (Watershed 3D) cell boundaries. The results shown here are from the demo dataset. We only show a part of the segmentation in the first six slices, because the original image size is too big (see Methods).

## Discussion

Automatic segmentation of 3D cellular images is a challenging task. Instead of improving the automatic method, we created a semi-automatic solution called Seg2Link, which uses deep learning predictions as input and assists users in quickly transforming imprecise predictions into a final accurate segmentation by providing an easy-to-use GUI as well as rich and convenient correction functions. Users can easily perform the first round of corrections with the Seg2D+Link module to resolve most errors within each slice, and then correct the remaining errors in the entire 3D image with the 3D correction module. Furthermore, we optimized Seg2Link’s data structure and algorithm to speed up correction and caching/saving operations, allowing users to efficiently segment large 3D cell images with billions of voxels.

Our software includes a number of useful functions for inspecting and correcting segmentation results, such as cell sorting/localization, division-relink, and undo/redo functions. Previously, some software programs have been developed to help users perform these automatic segmentation/manual corrections (Table 1). However, many only provide automatic segmentation (Berg et al. 2019, Ershov et al. 2021) or manual annotation/correction (Haehn et al. 2014; Zhao et al. 2018; Berger et al. 2018) without integrating them, making them less convenient to use. In addition, they often lack essential features required for efficient semi-automatic segmentation, such as masking regions outside the region of interest, automatic relinking of cells after a manual correction, multi-step undo/redo, fast caching/saving, cell sorting based on size, fast cell localization, and so on. Seg2Link adds those key features and simplifies the operations that users need to perform. For example, due to the imperfect predictions of cell boundaries, neighboring cells may frequently merge into one. In such cases, users must divide the cell into multiple pieces and relink the divided cells across slices, which requires numerous operations. With Seg2Link, users can easily complete these operations (division-relink) by pressing a single hotkey (Fig. 2C). The multi-step undo and redo feature is also essential, because misjudgments by users are common due to the difficulty of segmentation in real images. Our undo/redo function allows users to return to previous states, which greatly improves correction efficiency (Some of the other software programs do not implement this critical function. See Table 1). Also, saving intermediate results instantly after each operation is a must-have feature to avoid accidental loss. Our Seg2D+Link module employs a specific data structure to reduce the data size to be stored, allowing for fast data saving (Supplementary Fig. 2 and 5). Finally, users may only need to segment sub-regions or large cells. Our masking, cell sorting, and cell localization functions help users focus on segmenting cells of interest, greatly increasing efficiency (Fig. 2D and Fig. 3B). We did not include the function for predicting cell/non-cell regions in Seg2Link because the complicated deep learning training/predicting processes are better performed using other excellent programs (We provide a program using 2D U-Net, see Methods).

In addition to its rich functions, Seg2Link has been intensively optimized for processing 3D images in automatic segmentation, manual correction, and caching/saving. First, we designed a specific data structure to speed up the modification and data caching/saving in the Seg2D+Link module (Supplementary Fig. 2). We also designed a cache of bounding boxes to speed up the cell localization in the 3D correction module (Supplementary Fig. 4). Second, we tested the bottlenecks of our programs with profiling tools and optimized the time-consuming parts. Finally, all reference images (such as the raw image, prediction image) except for the segmentation results get loaded dynamically, remarkably reducing memory usage.

Apart from our excellent design for performing quick and easy segmentation/correction, the advantages of Seg2Link are also drawn from existing data processing tools developed by the scientific computing/image processing communities. These tools allowed us to write a concise program implementing both the upper-level visualization and the underlying computation. The recently released napari library, for example, includes a GUI for viewing and editing various types of image stacks. It also allows us to add new widgets and hotkeys to perform custom functions. NumPy, SciPy, and scikit-image are array/image processing libraries that allow us to write image processing functions. Dask, Python’s big data processing library, enables our software to dynamically load files on disk, reducing memory usage.

While Seg2Link is highly efficient in processing 3D images with billions of voxels, it has some limitations when processing even larger 3D images. First, the 3D correction module cannot process 3D segmentation exceeding the memory capacity (1 billion voxels in a 16-bit image roughly occupies 2 GB memory), which is a desired feature available in some other software for analyzing very large datasets such as entire brains (Berger et al. 2018; Zhao et al. 2018). This issue may be solved in the future by adding additional functions allowing users to divide the entire image into smaller sub-images to segment them separately, and then combine the results later. Second, Seg2Link currently only allows users to process and store the segmentation result as an 8-bit or 16-bit array, in order to save memory/disk space. As a result, users cannot handle images with more than 65,535 cells. This could be addressed in the future by allowing a 32-bit array.

Despite its limitations, Seg2Link can easily generate automatic segmentation using deep learning predictions and is superior in assisting users to efficiently inspect and correct errors. We believe Seg2Link is widely applicable to scientists analyzing 3D biomedical cell images in various types of studies, especially morphology and connectivity in the brain.

## Methods

### Computational environment

Seg2Link is entirely CPU-based and does not require a GPU. All of the analyses about the runtime in this manuscript were carried out on a laptop computer running Windows 11 with the CPU AMD Ryzen 9 5900HS and 32GB RAM. We also confirmed that the software can run on other desktop/laptop computers with Windows, macOS, or Linux systems. Seg2Link relies on napari, Dask, NumPy, SciPy, scikit-image, and other Python libraries for visualization and underlying computation. Users can easily install Seg2Link and its dependencies using the pip command.

### Image dataset

To demonstrate the functions of our software, we used a portion (1024 × 1024 × 1200 voxels) of a 3D EM image dataset (minnie65_8×8×40) of the visual cortex in mouse brain, which is publicly available (https://bossdb.org/project/microns-minnie). The dataset contains raw images as well as segmentation results with proofreading, with an x-y plane resolution of 8 nm/pixel and steps of 40 nm between slices. In 12 of the 1200 slices (i.e. slices # 50, 150, …, 1150), we transformed the provided segmentations to cell/non-cell regions as the ground truth for training a 2D U-Net model. The trained 2D U-Net was then used to predict cell/non-cell regions across all 1200 slices of the 3D image. These predictions and raw images (stored as 2D TIFF image sequence) were then imported into Seg2Link to perform the semi-automatic segmentation. We also used a larger portion (2048 × 2048 × 1200 voxels) of the same dataset to test the localization time with and without using bbox.

### Architectures of Seg2Link

#### Module 1: Seg2D+Link – Automatic segmentation

The automatic segmentation part of the Seg2D+Link module processes each slice of the 3D image one by one (Fig. 1A). The processing of every single slice consists of two sequentially executed steps: 2D segmentation and cross-slice linking (Fig. 1B).

The 2D segmentation procedure uses distance transformation to convert each slice of cellular/noncellular predictions into a distance map, finds local maximums as seeds, and then applies 2D watershed (Beucher and Meyer, 1993) to segment the image in the current working slice into individual cells (Fig. 1B). To mitigate over-segmentation, Seg2D+Link applies a Gaussian blur to the converted distance map and then uses the h-maxima transform (Soille, 1999) to filter the multiple local maxima within the same cell. By default, Seg2Link segments the entire 2D image in each slice. When the user provides a mask image indicating the region of interest (ROI), the program calculates the proportion of each segmented cell falling within the ROI. Cells with this proportion less than a user-specified threshold (default value 0.8) are deleted automatically (Fig. 2D).

From the second slice, the program will automatically link the segmented cells to the previous slice. For each pair of cells in the two adjacent slices, the program computes the overlap coefficient (Vijaymeena et al. 2016): Suppose there is a cell X in slice *i* and a partially overlapping cell Y in slice *i-1*. Their overlap coefficient is calculated using the equation below:

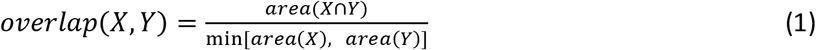

The calculated overlap coefficient is compared to a user-specified threshold (default is 0.5). When the overlap exceeds the threshold, the program will link X and Y to form a single cell (Fig. 1B).

#### Module 1: Seg2D+Link – Manual correction

Following the automatic segmentation and linking of each slice, the user can make the four types of corrections: 1. Merge multiple cells into a single cell; 2. Delete one or more cells; 3. Divide a cell in the current working slice into multiple cells; 4. Divide a cell in the current working slice and relink the results to the previous slices (Fig. 2C). Because of the specific data structure we used in the Seg2D+Link module to accelerate the modifications/caching/saving (see below), operations 1 and 2 can be applied to cells in any completed slice, whereas operations 3 and 4 can only be applied to cells in the current working slice. In module 2, 3D correction, users will be able to apply division and division-relink more freely in any slice (see below).

After executing each operation, our program will cache/save the current segmentation status in memory/hard disk. If users make mistakes, they can easily return to previous states using the undo/redo function (up to 10 steps by default, but users can modify it). If users quit the software, they can also easily restart from the point of interruption (but the previous states for undo/redo will be cleared).

Seg2Link makes use of the napari viewer to let users view and edit segmentation results (Fig. 2A). Users can pan and zoom the images to examine different regions in detail. The reference images and segmentation results can be overlaid to aid in visual inspection. The napari editing tools allow the user to select cells, and paint or correct their cell boundaries. In conjunction with our correction programs, these napari functions could assist users in quickly correcting the 3D segmentation results.

#### Module 1: Seg2D+Link – Underlying data structure

We designed a custom data structure for Seg2D+Link: The 3D segmentation result is saved as a series of 2D segmentation results corresponding to each slice. In each slice, the 2D segmentation is represented by a 2D label image (a 2D array) that stores the original 2D segmentation results from the 2D watershed, and a label list (a 1D list) that originally stores labels in the 2D segmentation result and will be updated when labels change due to link/merge/delete, etc. (supplementary Fig. 2). With this design, the program only needs to modify/cache/store a much smaller data structure: a 2D array + a group of 1D lists, rather than the large 3D array (supplementary Fig. 2). As a result, we are able to perform the modification and cache/save the intermediate state at a faster rate while using less memory/disk space.

While this custom data structure speeds up the functions, it requires an additional calculation process to display the updated segmentation results based on the original 2D segmentation results and the updated label lists. When there are too many slices to display, the computation becomes time-consuming. To reduce computational load, Seg2Link only displays segmentation results in a limited number of slices surrounding the current working slice (by default, 100 slices).

#### Module 2: 3D correction – Manual correction

The segmentation results from module 1 can be saved as a 3D array (npy format) and imported into module 2 to correct any remaining errors. The 3D segmentation results (2D TIFF image sequence) exported from other software can also be imported into module 2 for correction (e.g., The watershed 3D segmentation results shown in Fig. 4 were obtained using a different program and imported to the Module 3D correction). After completing all necessary corrections in module 2, the user can export the final segmentation results as 2D TIFF image sequence for further analysis in other software (Fig. 1A).

Module 2 offers the same four manual corrections as module 1: merge, delete, division, and division-relink. In contrast to module 1, the division and division-relink functions in this module can be conveniently implemented at any slice, but at the cost of a slower rate for modifications/caching/saving and increased memory/disk space requirements (See below).

In addition to the division and division-relink functions which are applied to a 2D subregion of a cell in a specific slice, we add a 3D division function in module 2 for dividing a cell in 3D space. We also add a function to insert a new cell that was not detected by deep learning, requiring users to manually paint the cell region with an automatically assigned new label (Fig. 3C).

#### Module 2: 3D correction – Easy inspection

Module 2 was designed to allow users to easily inspect the segmentation results. It allows users to do the following: 1. Sort cell ID numbers by size in descending order, so that large cells that may be more important can be easily selected and checked. 2. Remove cells smaller than a user-defined threshold, so that cells that are potentially irrelevant can be ignored. 3. Select a cell ID and jump to the central slice of the cell, which is useful when searching a cell through thousands of slices (Fig. 3B).

#### Module 2: 3D correction – Underlying data structure

In module 2, it is necessary to display the entire 3D segmentation results so that users can perform comprehensive visual inspections. The data structure in module 1 is no longer appropriate because it requires additional computations to display the updated segmentation results, which is time-consuming now. Instead, we use a simple data structure, a 3D array, to store the segmentation results (Supplementary Fig. 4).

One critical issue about using a 3D array as data structure is that searching for a cell’s location takes a long time due to the large search space. Seg2Link solved this problem by pre-calculating and storing the bounding boxes (bbox) of all cells in memory (Supplementary Fig. 4). Following each correction operation, the bbox of relevant cells are updated. In this way, the program can search for a cell in a much smaller sub-region and with much less time.

Another issue is that caching the entire 3D array is very space-intensive, making the undo/redo functions impractical. Our program solved this problem by caching changes only in the sub-region with modifications, which requires much less memory (up to 5 steps of undo/redo is possible by default, but users can modify this).

## Data availability

The Seg2Link source code and user guide for installing and using the software can be found at: https://github.com/WenChentao/Seg2Link. We have also provided a program for training a 2D U-Net to predict cell/non-cell regions, which can be found at: https://github.com/WenChentao/seg2link_unet2d. In addition, the demo dataset, including raw images and cell/non-cell predictions by the trained 2D U-Net, is available for download at: https://osf.io/wngty/.

## Author contributions

C.W., M.M., M.S., K.S. and K.D.K. designed the study. C.W. designed the architecture and wrote the software. C.W. and M.M. tested its functions, C.W. and K.D.K. wrote the manuscript. All of the authors have reviewed the manuscript.

## Acknowledgment

We are grateful to Nobuhiko Ohno, Shuichi Onami, Yusuke Azuma, and Koji Kyoda for discussion and technical advice. This work was supported by research grants from Japan Society for the Promotion of Science (JSPS) KAKENHI (20H05700 [to K.S. and K.D.K.]), Japan Agency for Medical Research and Development (AMED) (22gm1210007 [to K.S.]), Grant-in-Aid for Research at Nagoya City University (1921102 [to K.S. and K.D.K.]) and the Special Postdoctoral Researchers Program in RIKEN (to C.W.).

