## Supplementary figures for "Seg2Link: an efficient and versatile solution for semi-automatic cell segmentation in 3D image stacks"

### Supplementary material

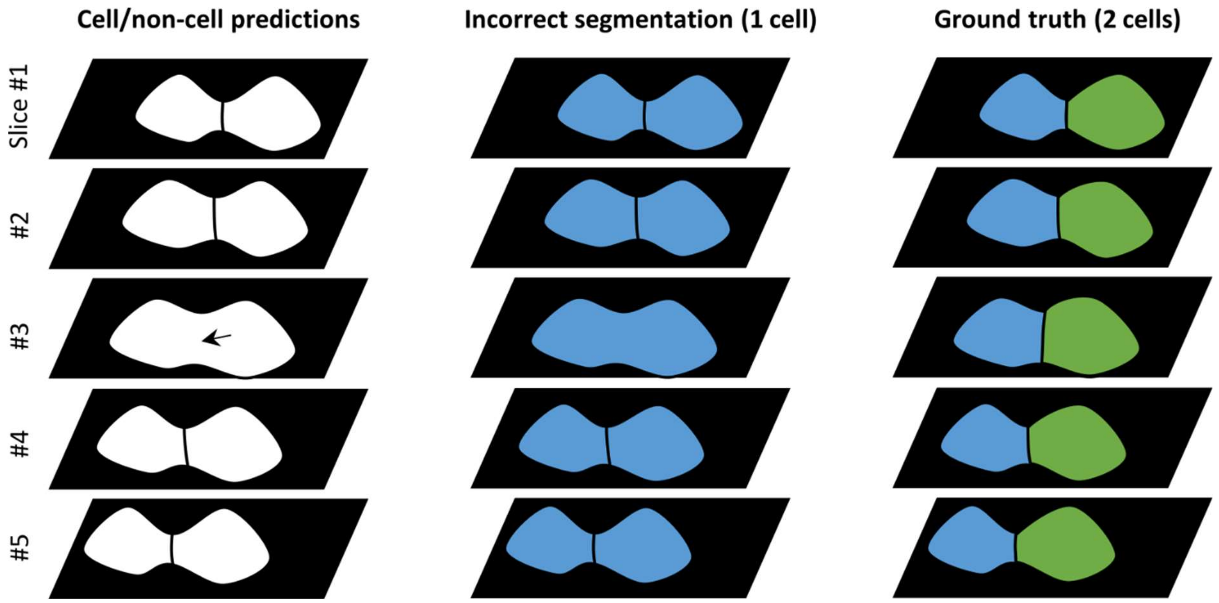


**Supplementary Figure 1**. An illustration of the effects of a local error (arrow) in cell/non-cell predictions on global segmentation results in 3D space.

A

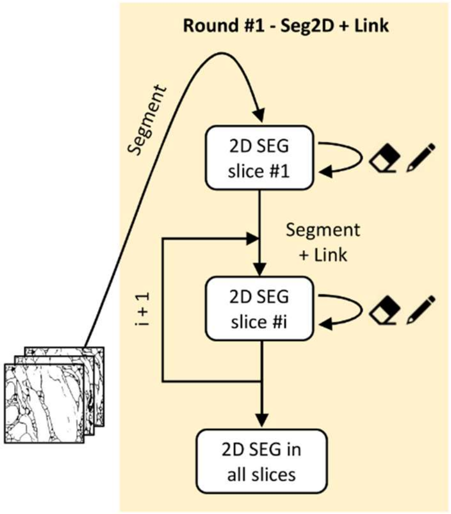


B

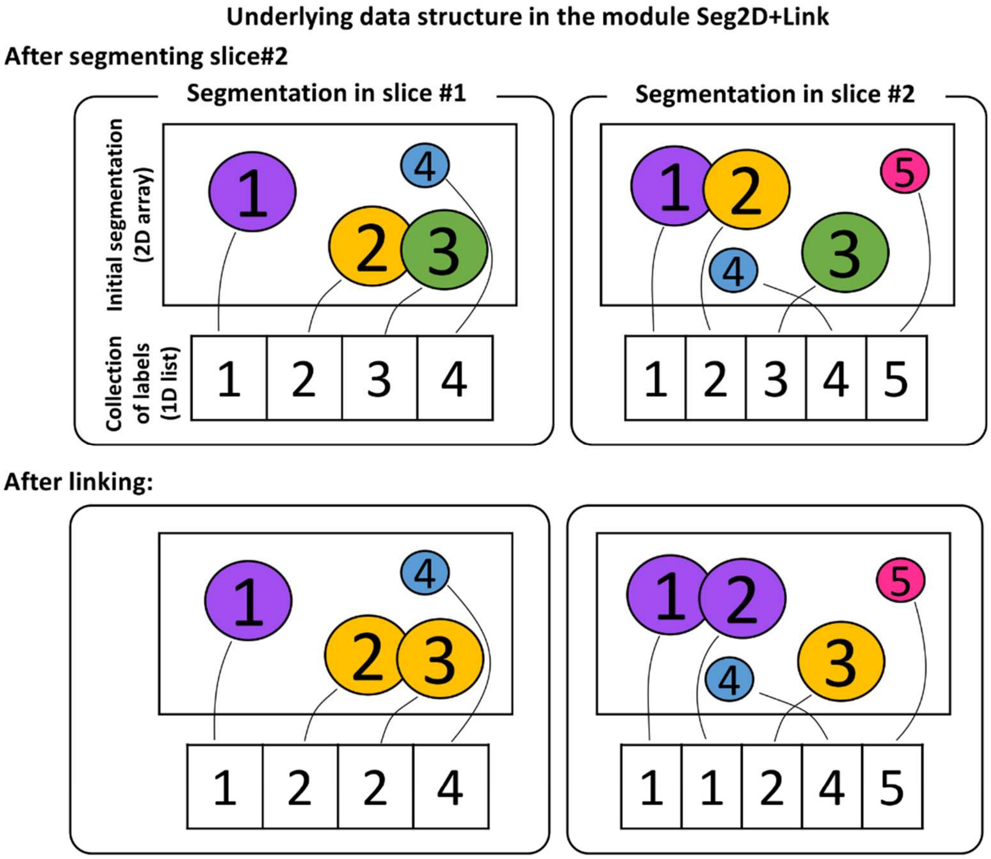


C

| Operations | Elements affected by the operation |
|---|---|
| Next slice (segment, segment + link) | 2D segmentation (c.w.s) + Label lists |
| Division/Division-Relink | 2D segmentation (c.w.s) + Label lists |
| Cache/Save intermediate state | 2D segmentation (c.w.s) + Label lists |
| Merge | Label lists |
| Delete | Label lists |

**Supplementary Figure 2**. The underlying data structure of the Seg2D+Link module. (A) The workflow in the Seg2D+Link module. (B) A diagram of the underlying data structure, as well as the overlap linking process based on the data structure. (C) Operations that alter elements of the data structure. c.w.s: current working slice.

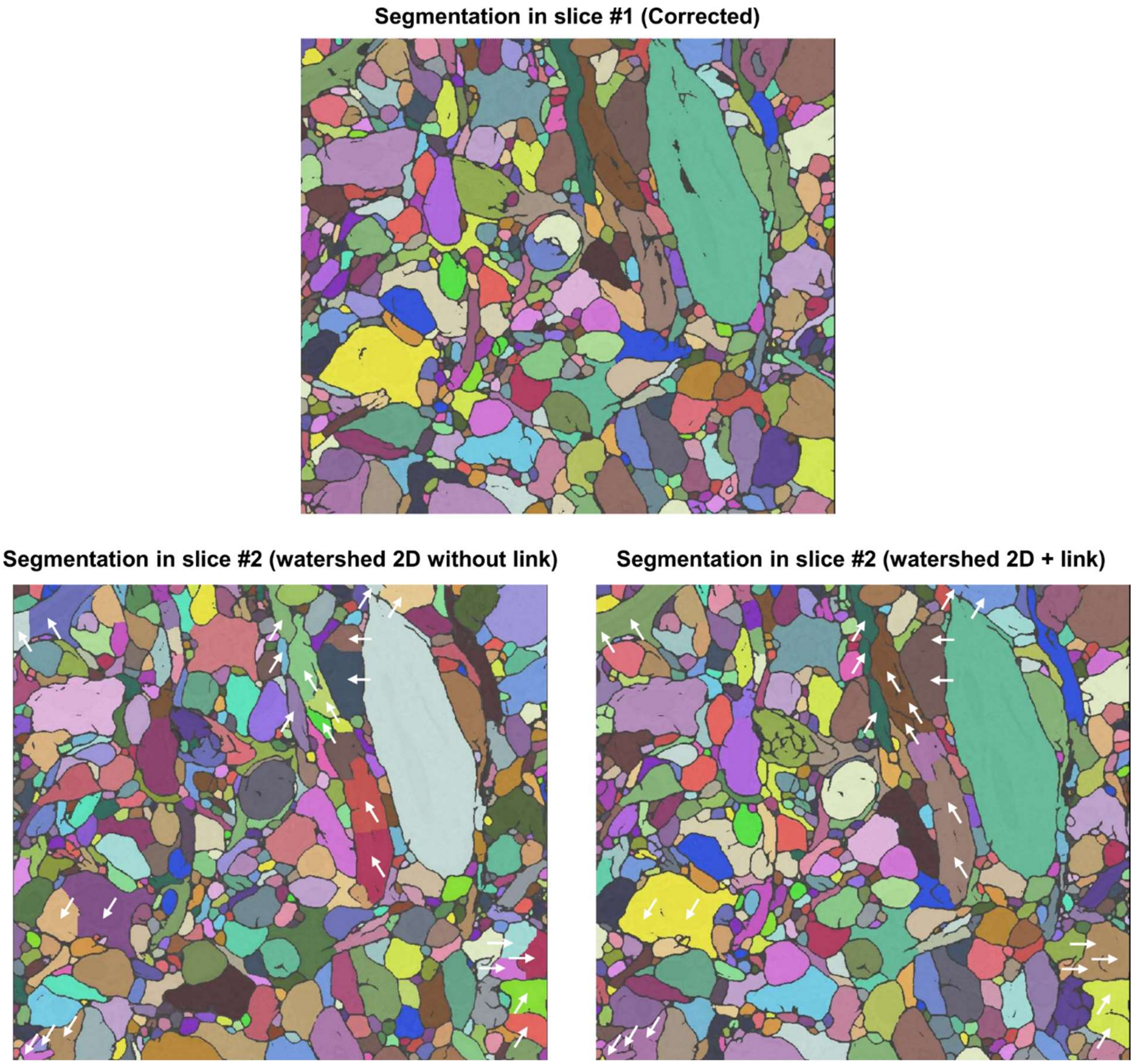


**Supplementary Figure 3**. Corrections made in a previous slice can improve automatic segmentation in subsequent slices. (Top) Manually corrected segmentation in slice #1. (Bottom left) Automatic segmentation in slice #2 using watershed 2D but no link. (Bottom right) Automatic segmentation in slice #2 using watershed 2D + link. The arrows point to the same regions that were split incorrectly by watershed 2D and correctly merged by watershed 2D + link. Colors were automatically assigned by napari.

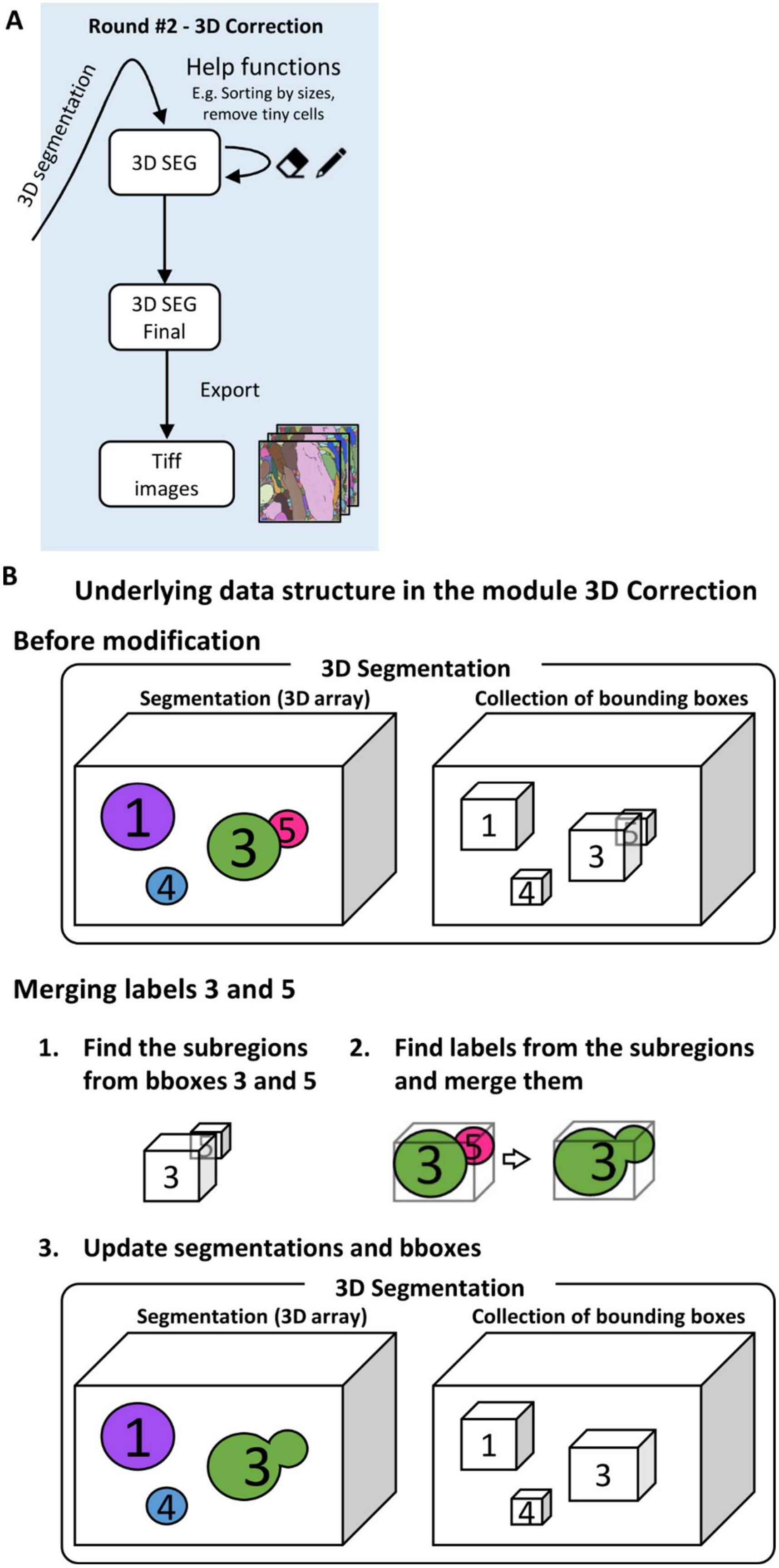
A
Round #2 - 3D Correction
3D segmentation
Help functions
E.g. Sorting by sizes, remove tiny cells
3D SEG
3D SEG Final
Export
Tiff images
B
Underlying data structure in the module 3D Correction
Before modification
3D Segmentation
Segmentation (3D array)
Collection of bounding boxes
Merging labels 3 and 5
1. Find the subregions from bboxes 3 and 5
2. Find labels from the subregions and merge them
3. Update segmentations and bboxes
3D Segmentation
Segmentation (3D array)
Collection of bounding boxes

**Supplementary Figure 4**. The underlying data structure of the 3D correction module. (A) The workflow in the 3D correction module. (B) A diagram of the underlying data structure, as well as the merge process based on the data structure.

**A**

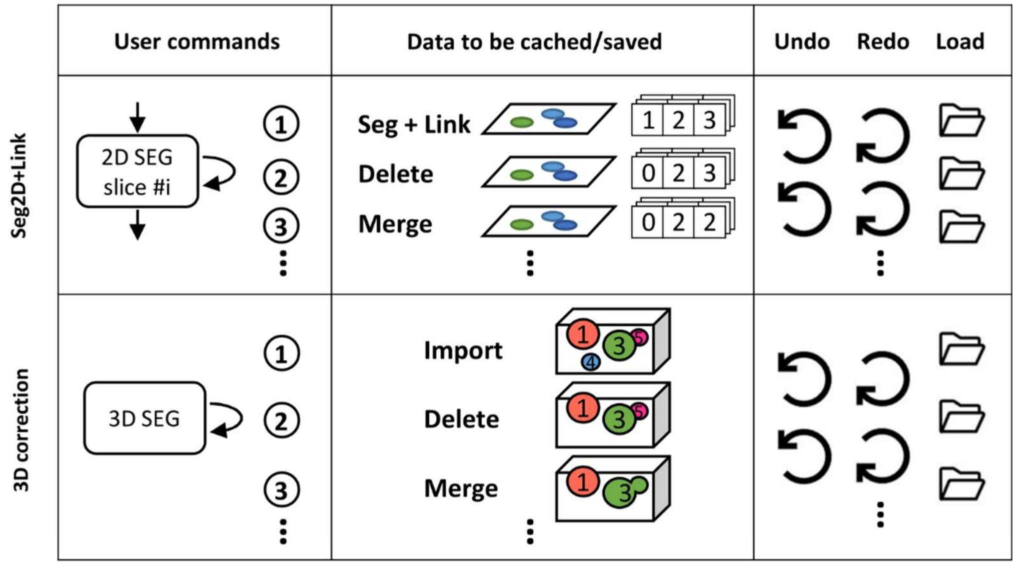


**B**

| Module | Data to be saved | Data size | Time for saving |
|---|---|---|---|
| Seg2D+Link | 2D array (c.w.s)<br>+ label lists (1 – c.w.s) | **Total:** 82 KB ~ 2.0 MB<br>- 2D array: 80 KB / slice<br>- Label list: 1.61 KB / slice | 0.8 ~ 20 ms |
| 3D correction | 3D array | **Total:** 2.46 GB | 25 sec |

**Supplementary Figure 5**. The caching/saving methods used in the Seg2D+Link and 3D correction modules. (A) The data to be cached/saved in the two modules after performing each user command. Note that in reality, our 3D correction module only caches a subregion of the entire 3D array to reduce the memory utilization. (B) A comparison of the two modules' efficiency in saving a real data (demo dataset). Our software saves each 2D array in npz format (compressed), label lists in pickle format, and 3D arrays in npy format (without compression since it's time-consuming for large data). The time is estimated assuming the write speed of the hard disk is 100 MB/sec. c.w.s: current working slice.
